# A Required Sensitivity Analysis for Predictive Reliability of Genome–Scale Metabolic Networks: Probing Biomass, Nutrient Conditions, and Specialised Metabolism with *Penicillium rubens* as a Case Study

**DOI:** 10.1101/2025.09.10.675325

**Authors:** Nègre Delphine, Larhlimi Abdelhalim, Bertrand Samuel

## Abstract

The Biomass Objective Function is crucial to the predictive capability of Genome–Scale Metabolic Network (GSMN) models. Its definition should theoretically reflect the organism-specific macromolecular composition and stoichiometry under defined environmental conditions. In practice, however, reconstruction decisions—whether documented, reasoned, or arbitrary—often lead to model refinements which can potentially introduce ambiguities and may compromise the reliability of simulation outcomes. To mitigate this issue, we propose that systematic sensitivity analysis should be a mandatory step in GSMN validation. This approach quantitatively assesses the reliability of flux predictions by probing a model’s responsiveness to perturbations in its core objective function and environmental inputs. We demonstrate this approach using the fungus model *Penicillium rubens i*Prub22. First, we evaluate the sensitivity of predicted fluxes to variations in the stoichiometric coefficients of the biomass reaction. Then, we examine the model’s metabolic behaviour under alternative nutrient conditions. Finally, we assess whether secondary metabolite production, governed by its own regulatory logic, remains robust to changes in the biomass objective function formulated for growth. Together, these analyses measure the degree to which a model’s predictions are sensitive to specific reconstruction choices, thereby establishing a standard for evaluating predictive robustness to parameter uncertainties and functional quality in GSMNs.

## 1. Introduction

Since 2010, the reconstruction of Genome–Scale Metabolic Networks (GSMNs) has been guided by a reference protocol (Thiele & Palsson, 2010). However, the diversity of metabolic databases, coupled with the wide range of reconstruction and analysis tools, and the inconsistent definition and application of standards (Carey et al., 2020; Ebrahim et al., 2015) have often led to models functioning as “black boxes”, with key reconstruction decisions insufficiently documented. This lack of transparency limits their reusability and complicates the assessment of their predictive accuracy, as the reproducibility of results remains a notable obstacle (Tiwari et al., 2021).

This naturally raises the question of what constitutes GSMN quality. Intuitively, the quality of a reconstruction refers to the reliability of the underlying data and the robustness of the methods used in reconstruction (Heirendt et al., 2019; Hucka et al., 2018; Moretti et al., 2021; Novère et al., 2005; Olivier & Bergmann, 2018). This encompasses the accuracy of gene and protein annotations, the correctness of the associated biochemical reactions, and the consistency of the experimental data integrated in the model. Furthermore, GSMN quality relies on accurately representing metabolite-enzyme interactions and the model’s ability to predict system-level biological behaviour across diverse conditions. Despite its importance, the question of how to evaluate GSMN quality remains ambiguous. To our knowledge, beyond the MEMOTE test suite, there are few established tools or measures specifically designed to assess the quality of published reconstructions and models. Moreover, existing assessments predominantly address standardisation, formatting and encoding compliance, thereby evaluating structural aspects rather than biological content. While such structural considerations are essential for maintenance and sharing, the core value of a GSMN lies in its capacity to model the biological phenomena of the organism it represents. In practice, however, the evaluation of this capacity is often reduced to a single criterion: the model’s ability to simulate organismal growth — typically through a functional biomass reaction.

Traditional analyses of GSMNs typically rely on flux-based approaches such as flux balance analysis (FBA) (Orth et al., 2010) and flux variability analysis (FVA) (Gudmundsson & Thiele, 2010). These methods are applied under defined conditions and assume the system operates at steady state. The problem is formulated mathematically as a linear optimisation, where the objective function generally represents organism growth. Flux balance analysis aims to predict a single optimal flux distribution that maximises the objective, while FVA explores the range of feasible fluxes for each reaction that is consistent with this optimum. These approaches are widely used to study metabolic capabilities, identify essential reactions, and explore and delineate the solution space compatible with physiological steady-state constraints.

The growth simulation relies on the formulation of the biomass objective function (BOF), an artificial construct representing the sum of macromolecules required for cellular structure and the energy necessary for their synthesis (Feist & Palsson, 2010). The BOF, thus, reflects the dynamic interplay between catabolic pathways, releasing energy and precursors, and anabolic pathways that utilise these for biosynthesis and maintenance. Biomolecules, mainly carbon-based, form the cellular framework, with functional properties defined by specific groups. Carbohydrates, lipids, proteins, and nucleic acids constitute the central macromolecular classes, most of which are polymers of monomers. Accurate modelling of metabolism and growth requires a detailed understanding of their composition and functions.

Once the nature of these biomolecules is established, their quantities—as molar fractions and thus expressed in the stoichiometric coefficients of the BOF—must be determined. Ideally, they are based on experimentally measured data from defined culture conditions (*e.g.,* continuous growth in a chemostat), with coefficients normalised per gram of dry biomass (Thiele & Palsson, 2010). This detailed characterisation enables precise calculation of biomass yields from stoichiometry. Beyond quantitative aspects, the qualitative composition of the biomass reaction—*i.e.,* the inclusion or exclusion of specific precursors—is critical for *in silico* gene knock-out simulations and identifying essential genes and reactions. In recent years, despite significant progress in improving the accuracy of BOF definitions (Lachance et al., 2019), the need to standardise the acquisition and integration of this reaction persists (Chan et al., 2017).

According to this definition, and as expected, the BOF is highly specific to the organism under study. Nevertheless, its formulation is often inherited from similar species due to the experimental challenges associated with its accurate determination (Beck et al., 2018). This aspect represents one of the main sources of uncertainty in defining biomass composition, an issue that is often underestimated (Bernstein et al., 2021). Indeed, biomass reactions from more or less related organisms are regularly used to characterise the organism of interest without necessarily considering the variations between different species (Xavier et al., 2017). When the BOF employed in a model is derived from another reconstruction—whether from the same organism or a related species—a quality check of the biomass reaction can be performed using sensitivity analyses.

Beyond differences in biomass composition between organisms, variability may also exist within the same species. Changes in growth rate, nutrient availability, or physical parameters can affect organism growth (Bernstein et al., 2021), and cells respond through metabolic adjustments that influence the biosynthesis of metabolic precursors required for building cellular components. Therefore, it is essential to consider the dependency of biomass composition on environmental conditions (Schulz et al., 2021).

The fundamental objective of sensitivity analyses is to assess model robustness and identify critical parameters (Borgonovo & Plischke, 2016; Nobile et al., 2021). These analyses examine how an optimal output state of a mathematical model responds, qualitatively or quantitatively, to various changes in model inputs. For example, sensitivity analysis of the BOF to variations in biomass precursor coefficients (Nanda et al., 2020) helps verify whether *in silico* growth is sensitive to all biomass precursors. Similarly, sensitivity analysis of exchange reaction boundary variations (Lachance et al., 2021) allows testing model responses to environmental changes and, where relevant, determining optimal ranges for designing different culture media.

Therefore, we advocate for the systematic application of three complementary sensitivity analyses to evaluate the robustness and predictive reliability of GSMN through the BOF definition. First, we perturb the stoichiometric coefficients of the biomass reaction to assess the extent to which simulated growth depends on its quantitative formulation, thereby testing the robustness of biomass definition to parameter uncertainty. In a second step, we simulate growth across a spectrum of nutrient conditions to evaluate how biomass production responds to environmental constraints encoded in the model, and to identify potential limitations or artefacts arising from media definition. Finally, once the sensitivity of growth to biomass formulation has been characterised, once again through quantitative variations in the biomass reaction, we examine their impact on metabolic outputs that are potentially not directly coupled to the growth objective, using specialised metabolite production as a representative case. Together, these analyses provide tractable means to evaluate the influence of the BOF definition and environmental conditions on a model’s predictive fidelity.

To illustrate this approach, we selected the *Penicillium rubens* model *i*Prub22 (Nègre et al., 2023), the GSMN of a filamentous fungus which exhibits substantial biosynthetic capacity for specialised metabolism production (Eagan & Keller, 2025), allowing us to test predictions beyond core growth. This parameterised reconstruction has a MEMOTE quality score of 74%, a metric which evaluates the structural quality of *i*Prub22 but provides no insight into predictive performance (Lieven et al., 2020).

Modelling specialised metabolism is a demanding task because these pathways are related to responses to biotic and/or abiotic factors that may explain specific phenotypes. In nature, these pathways are considered relatively unlinked to the organism’s growth objective (Casas López et al., 2004; Hope et al., 2005). However, some of these metabolites rely on molecular building blocks that are also common to the BOF, creating potential interactions between biomass definition and the predictions of specialised-metabolite production. Assessing how biomass coefficients influence these outputs is therefore essential for understanding how BOF definition may affect simulated specialised-metabolism phenotypes. Therefore, we also explore, for the first time, the use of sensitivity analysis, focusing on non-objective reactions to assess the influence of the BOF on the specialised metabolism. This evaluation bridges core metabolism, growth, and specialised metabolism, enabling a more comprehensive assessment of model performance in capturing physiologically relevant phenotypes. These simulations serve to evaluate the impact of biomass formulation on the independence of predictions and on model sensitivity, criteria that, when met, attest to the quality of the model used. Such insights will help advance understanding of the relationships between environment, metabolism, and biomass composition in the organism under study.

## 2. Materiels & Methods

### 2.1. Model and Computational Environment

The GSMN model used in this study corresponds to the most recent reconstruction of *P. rubens*, *i*Prub22 (Nègre et al., 2023). The model is publicly available *via* BioModels under the identifier MODEL2306150001.

All simulations were primarily conducted in MATLAB (MathWorks Inc., Natick, MA, USA—R2021a) using the COBRA Toolbox v3.4 (Heirendt et al., 2019) and the Gurobi optimiser (v12.0.2). Simulation results were exported to R (v4.3.3) and visualised in RStudio using the ggplot2 package (v3.4.2) (Wickham, 2016). Alternatively, simulations and visualisations can be reproduced using Python (v3.12.7) and COBRApy (v0.29.0) (Ebrahim et al., 2013), thereby avoiding reliance on proprietary software. All scripts required for both workflows (*i.e.,* MATLAB–R and Python-only) are archived on Zenodo (https://doi.org/10.5281/zenodo.17091974). To ensure full reproducibility, Conda environment files are supplied for both the R-based visualisation and the Python-based simulation and plotting pipelines. Interactive files are provided in their native formats—MATLAB LiveScripts, R Markdown, and Jupyter Notebooks—as well as in HTML-rendered versions to support transparency, reproducibility, and exportability. The MATLAB/R process requires approximately 22 min from model loading to data generation and plotting, whereas the Python-based pipeline takes approximately 26 min. All simulations were executed on a Dell Precision 3591 mobile workstation running Ubuntu 22.04.5 LTS (Linux kernel 6.8.0-60-generic, x86_64 architecture). The machine was equipped with an Intel® Core™ Ultra 7 165H processor (16 physical and 22 logical cores) and 66.85 GB of RAM.

### 2.2. Constraint Specification

The default model constraints were maintained. The minimal nutrient set required for model functionality included a carbon source, a nitrogen source, molecular oxygen (Uptake_136), ferrous iron (Uptake_062), sulphur (Uptake_169), inorganic phosphate (Uptake_146), thiamine (Uptake_171), and riboflavin (Uptake_157). The latter two were necessary to activate the cofactor (COF) subsystem by enabling the biosynthesis of thiamine diphosphate and flavin adenine dinucleotide (FAD). It should be noted that flavin mononucleotide (Uptake_065) and hydrogen peroxide (Uptake_077) can be used as substitutes for riboflavin and the oxygen source, respectively. Oxygen uptake was left unconstrained. The upper bounds for carbon and nitrogen sources were 15 and 5 mmol.gDW⁻¹.h⁻¹, respectively. All other mentioned uptakes were left to 10 mmol.gDW⁻¹.h⁻¹.

Uniquely, for specialised metabolite production analyses, three additional uptake reactions were enabled to account for cofactor and precursor requirements: Uptake_155 (Red-NADPH-Hemoprotein-Reductases, *i.e.,* a class of enzymes involved in redox reactions using NADPH to reduce haemoproteins), Uptake_044 (decanoate), and Uptake_144 (phenoxyacetate). Each was constrained to 10 mmol.gDW⁻¹.h⁻¹.

### 2.3. Stoichiometric Sensitivity of the Biomass Reaction

To assess the model’s sensitivity to quantitative variations in the biomass reaction (Biomass_rxn), the stoichiometric coefficient of each reactant in the biomass equation, except for ATP, was independently modified by ±50% of its original value. To refine the resolution of the analysis, the same perturbation strategy was subsequently applied to the seven metabolic subsystems responsible for the biosynthesis of each biomass precursor: cofactors (r1465), DNA (r1458), RNA (r1457), phospholipids (r1460), cell wall components (r1455), proteins (r1456), and the free amino acid pool (r1459). In all FBA simulations, the biomass objective function was maximised at 100%.

For specialised metabolite production, FVA was performed under the constraint of 80% maximal biomass flux to estimate theoretical upper bounds for key precursors (*i.e.,* metabolic branch points) or final products of specialised metabolites known to be produced by *P. rubens*. The selected specialised metabolites encompass those documented in *P. rubens* Wisconsin 54-1255 and span the major biochemical classes of specialised metabolism (*i.e.*, non-ribosomal peptides, polyketides and terpenes) (Guzmán-Chávez et al., 2018; Martín et al., 2017; Van den Berg et al., 2008). Given the incomplete coverage of specialised metabolism in current databases, when necessary, we restricted our analysis to the last known metabolic branching points linked to an orphan specialised metabolite under investigation.

Only stoichiometric coefficients from the biomass reaction itself were tested. The relevant production reactions were those leading to the production of: isopenicillin N (Production_010) and 6-aminopenicillanate (Production_001) for penicillins; 3-methylphenol (Production_016) for isoepoxydon and yanuthones; histidyltryptophyldiketopiperazine (HTD) (Production_022) for roquefortines; YWA1 (Production_033) for melanin; aristolochene (Production_036) for PR-toxin; 3,5 dimethylorsellinate (Production_037) for andrastin; ferrichrome (Production_034) and ferrichrome A (Production_035); averufin (Production_038); and versiconol (Production_040).

### 2.4. Sensitivity of Biomass Yield to Nutrient Availability

Robustness analysis was performed using the robustnessAnalysis function in the COBRA Toolbox. This method evaluates the sensitivity of a target flux—here, biomass production—to variations in individual reaction—here, nutrient uptake fluxes (Becker et al., 2007). For each selected uptake reaction, the flux was fixed to 500 evenly spaced values between its minimum and maximum bounds. At each fixed value, FBA was conducted to maximise biomass production, and the resulting objective values were recorded. These data were exported to R CRAN and used to generate robustness profiles, representing biomass production as a function of nutrient availability. Carbon and nitrogen sources yielding strictly identical robustness profiles (*i.e.,* identical biomass values across all tested uptake rates) were grouped. A total of 31 carbon sources and 28 nitrogen sources (Table S1 in Appendix A) were tested as potential substitutes for glucose and ammonia. Ammonia served as the fixed nitrogen uptake during carbon source evaluations; glucose was used as the fixed carbon source during nitrogen source analyses. The values displayed by a vertical bar on the robustness plots (Figure 3.A) correspond to the nutrient uptake rates required to reach the organism’s theoretical and experimental growth rate, set at 0.20 h⁻¹ (Nielsen, 1997).

## 3. Results

### 3.1. Assessment of Growth Robustness under Biomass Objective Function Perturbations

To assess the BOF robustness of the GSMN *i*Prub22, we examined the growth’s sensitivity to variations in its stoichiometric coefficients and its subsystems (Figure 1).

**Figure 1.**
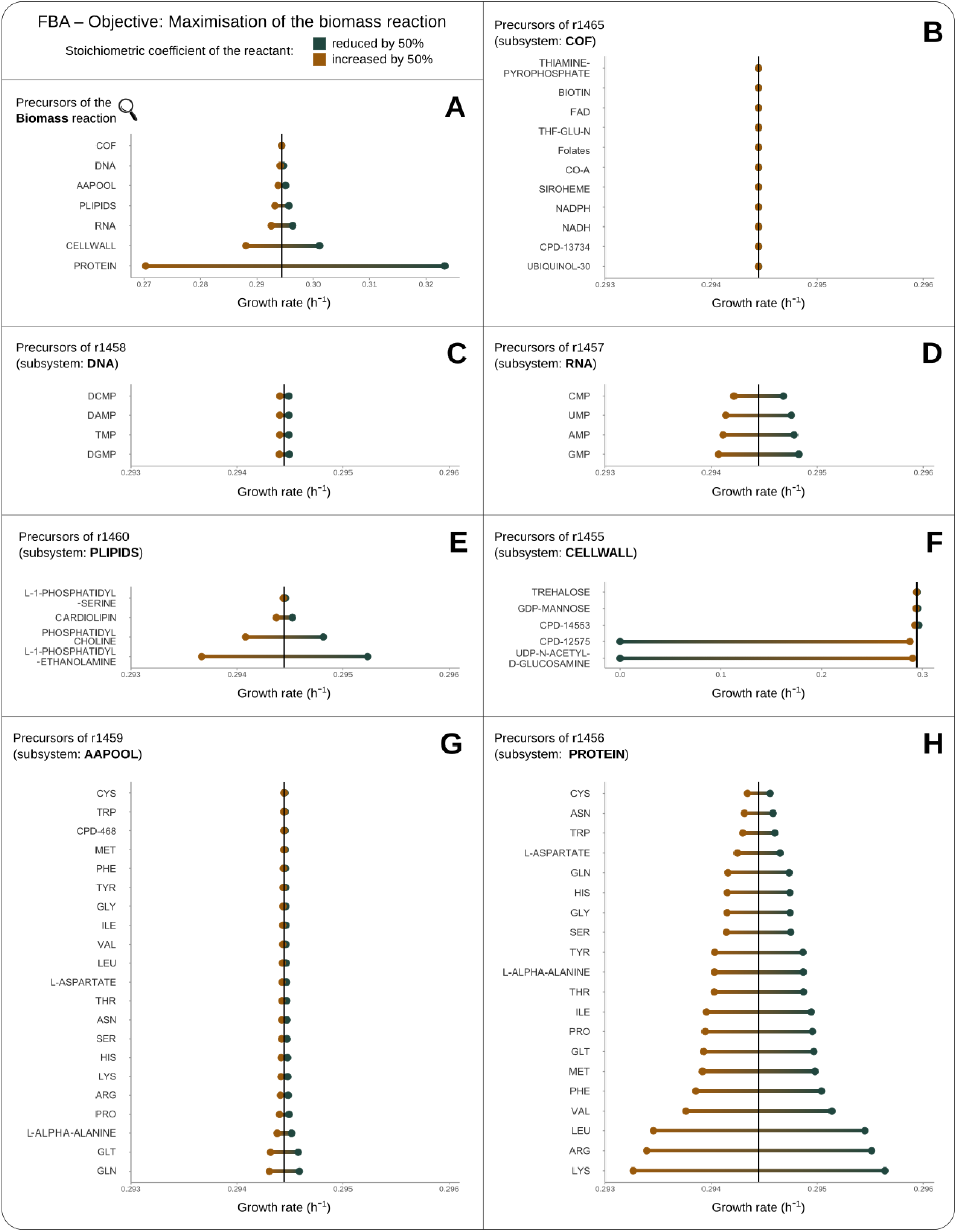
Sensitivity analysis of the biomass objective function to variations in the coefficients of biomass precursors (A) and its subsystems (B-H). FBA was maximised to 100% for each variation in the coefficient of the precursor considered. The model constraints correspond to the default settings. The scales of the panels were harmonised, except for panels A and F, which display variations of larger magnitude. Respectively, brown (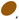) and green (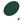) dots represent the FBA results for a 50% increase and decrease in the coefficient value. The black vertical line indicates the biomass flux under default conditions, i.e., 0.29 h⁻¹. Subsystem precursors are identified using their MetaCyc identifiers.

Each reactant within the BOF, and subsequently within each biomass subsystem, except for ATP (*i.e*., representing growth-associated maintenance or the energetic cost of polymerisation in the PROTEIN, DNA, and RNA subsystems), was individually perturbed by ±50% of its original stoichiometric coefficient. FBA was then used to estimate the new optimal predicted specific growth rate change. Under default conditions, the model predicts a growth rate of 0.29 h⁻¹.

Across the BOF components, the maximum observed variation in predicted growth rate was 0.06 h⁻¹ (Figure 1.A), indicating overall robustness. The most pronounced effects were associated with the PROTEIN and CELLWALL subsystems.

Within each synthetic precursor subsystem, the amplitude of the observed variations ranged from 10⁻³ to 10⁻⁶ h⁻¹. Given a solver feasibility tolerance of 10⁻⁶, no meaningful impact can be attributed to the unitary definition of coefficients within the cofactor subsystem (Figure 1.B). Variations observed for the DNA subsystem, on the order of 10⁻⁵ h⁻¹, are likewise negligible (Figure 1.C). The RNA (Figure 1.D), phospholipid (Figure 1.E), and free amino acid pool (Figure 1.G) subsystems displayed only minor effects, with amplitudes around 10⁻⁴ h⁻¹. The largest variations were observed for the PROTEIN subsystem (Figure 1.H), reaching up to 10⁻³ h⁻¹, suggesting a slightly increased sensitivity to the quantitative definition of specific amino acids.

However, two CELLWALL precursors—UDP-*α*-glucose (CPD-12575) and UDP-*N*-acetyl-*α*-D-glucosamine (UDP-N-ACETYL-D-GLUCOSAMINE)—yielded a predicted growth rate interval that does not include the model’s optimal reference value (Figure 1.F), suggesting potential vulnerabilities requiring further scrutiny. With the exception of these two elements, the range of variation between the decrease and the increase of each coefficient is symmetric around the model’s default predicted growth value and the predicted optimal growth rate of *i*Prub22 remains relatively robust with respect to quantitative variations in BOF composition.

### 3.2. Assessment of Growth Phenotypes under Nutrient Condition Perturbations

To evaluate the metabolic adaptability of *i*Prub22 to environmental changes, we examined how variations in the uptake fluxes of key nutrients affect biomass production. Using single variable robustness analysis, we characterised the system’s response to individual nutrient constraints, aiming to identify thresholds, sensitivities, and potential inhibitory effect (Figure 2.A). This approach provides insight into the physiological relevance of nutrient availability and the functional limitations embedded in the network structure.

**Figure 2.**
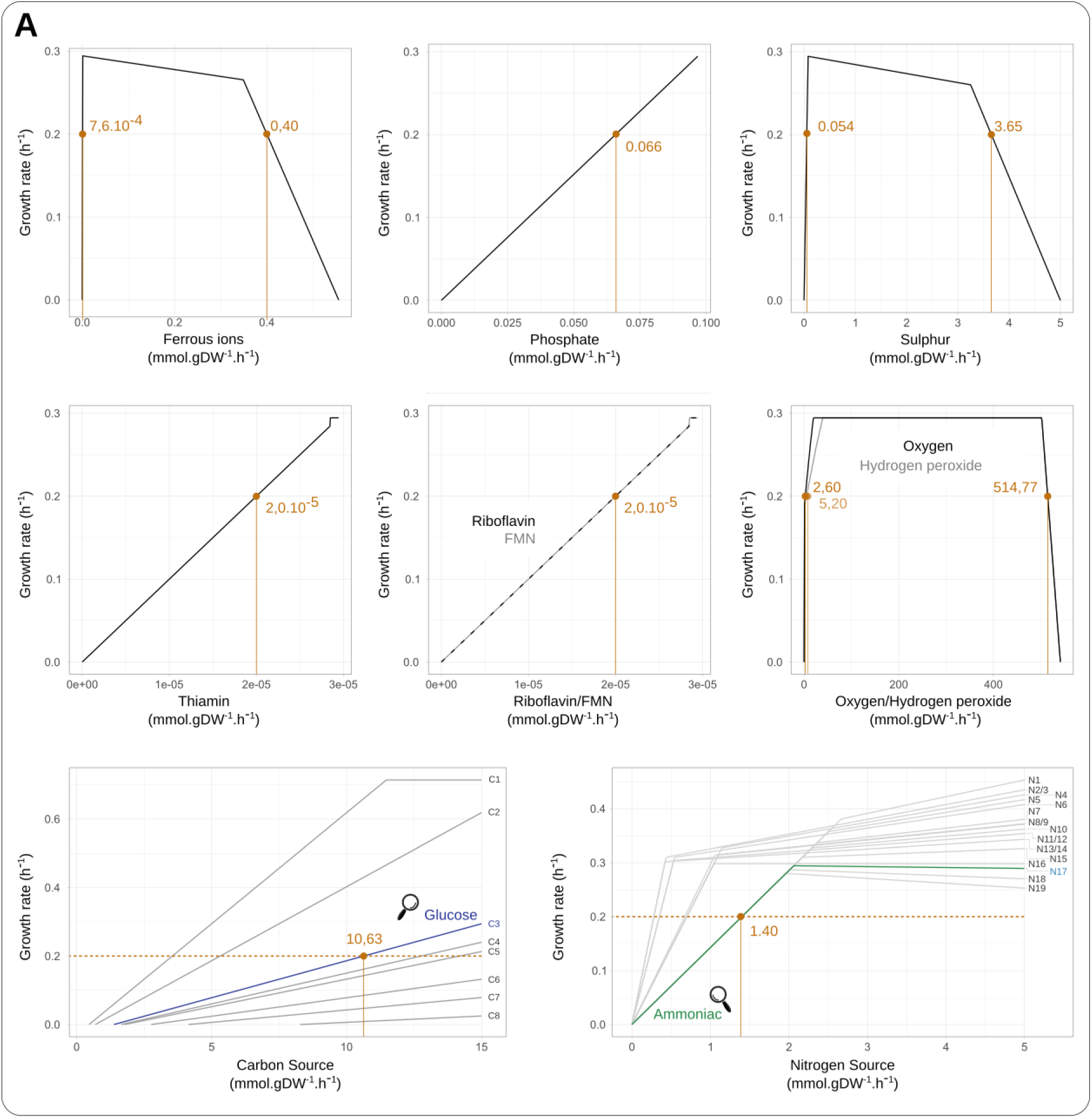
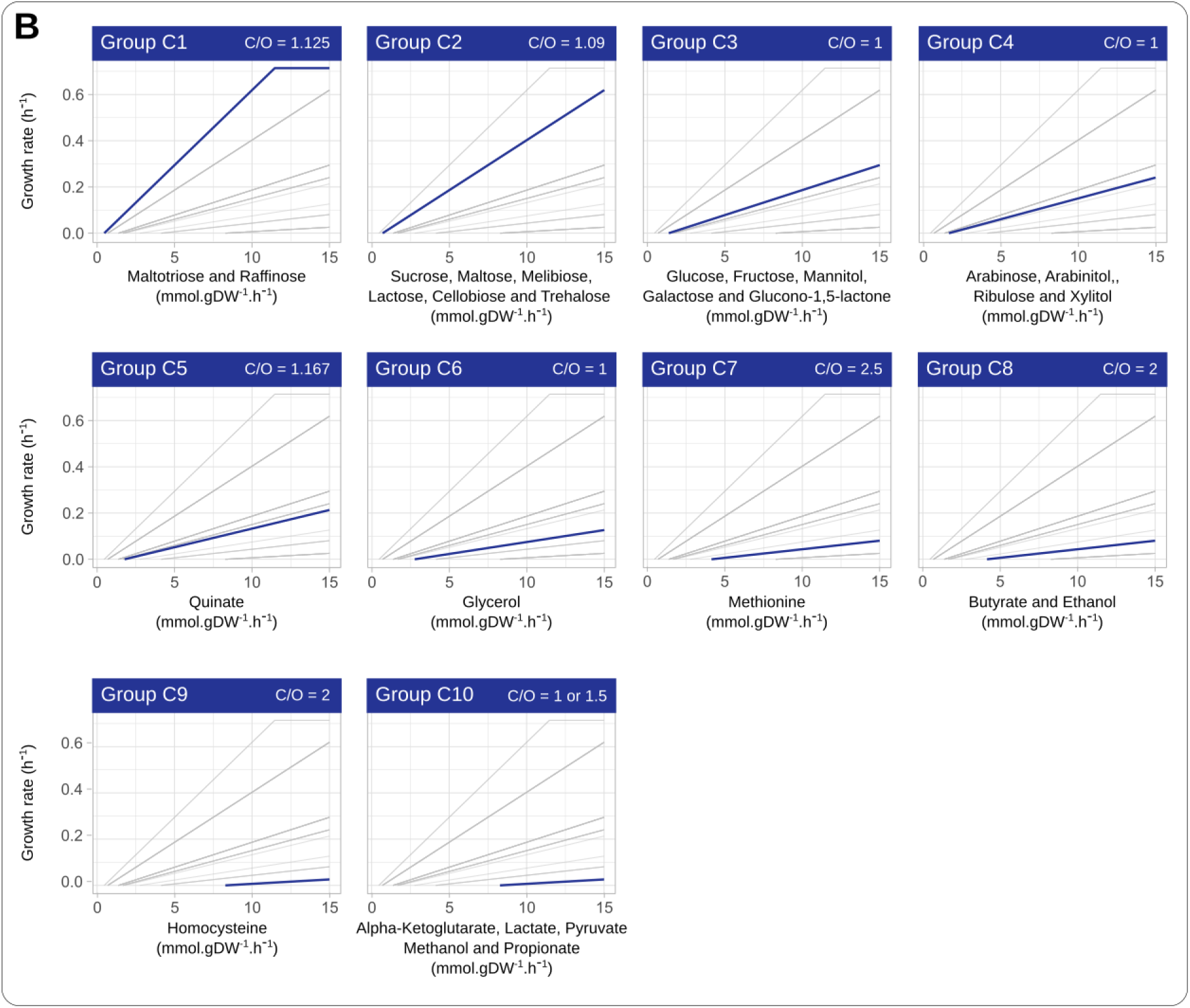
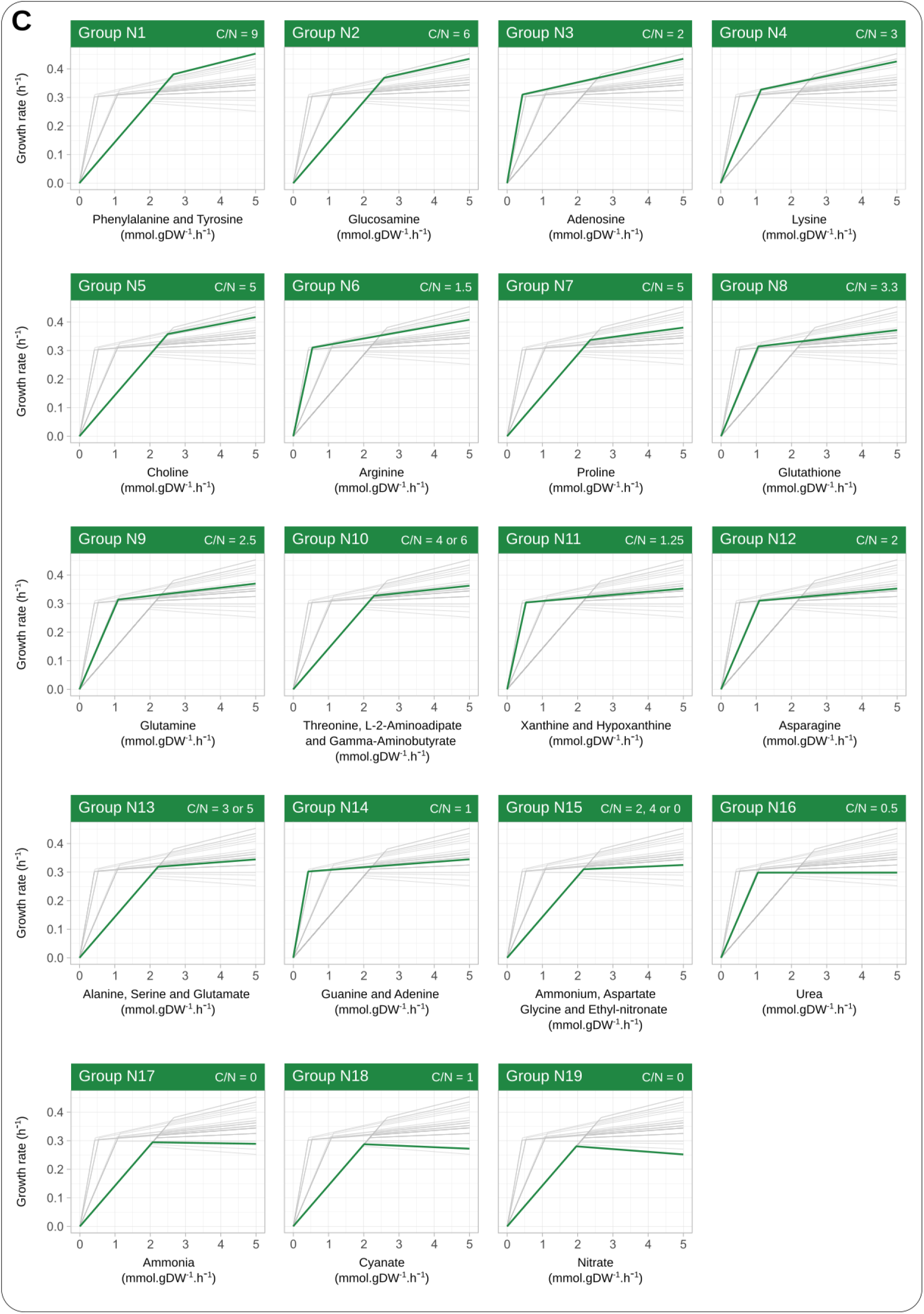
Robustness analyses of the required uptake rates for the functionality of iPrub22. (A) Robustness profiles for the minimum import fluxes are listed in the Materials & Methods section. Orange dots indicate the import flux values required to reach a biomass production rate of 0.20 h⁻¹, corresponding to a classical experimentally determined growth rate (Nielsen, 1997). (B) Sensitivity analysis of various carbon sources, grouped into eight categories according to carbon content (C1-C8). Except for groups C5, C7, and C8, the results show that a lower carbon-to-oxygen ratio (i.e., lower carbon availability) and a lower carbon number generally require higher uptake rates to reach the target biomass value. The imposed upper limit of 15 mmol.gDW⁻¹.h⁻¹ does not allow saturation plateaus to be reached. (C) Sensitivity analysis of various nitrogen sources, grouped into nineteen categories according to nitrogen content (N1-N19). For nitrogen sources, the carbon-to-nitrogen ratio appears to have a less pronounced effect on biomass production than the carbon-to-oxygen ratio observed for carbon sources.

**Figure 3.**
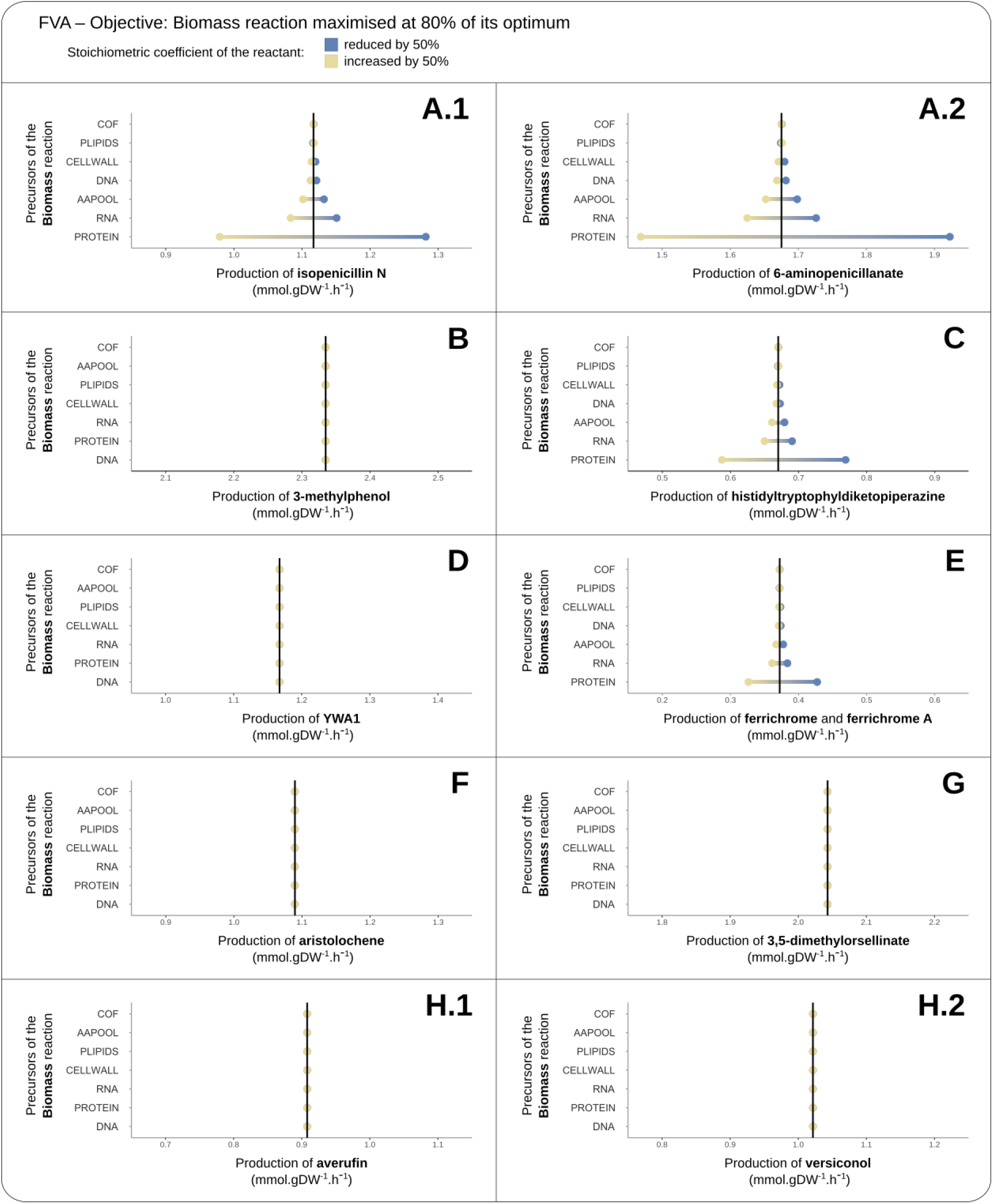
Sensitivity analysis of specialised metabolite production fluxes in Penicillium rubens in response to variations in biomass precursor coefficients. An FVA maximising the biomass reaction to 80% was performed for each variation in the coefficient of a biomass precursor. Yellow (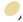) and blue (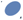) dots represent the maximum FVA values obtained after a 50% increase and decrease, respectively, in the coefficient of each precursor. Reactants are ranked by decreasing amplitude, with scales standardised across panels. The selected precursors (i.e., branching points) are representative of the main biosynthetic pathways for (A) penicillins, (B) isoepoxydon and yanuthones, (C) roquefortines, (D) melanin, (E) ferrichromes, (F) PR-toxin, (G) andrastins, and (H) averufin and versicolorin B. The black vertical line represents the maximum production flux under the default biomass precursor coefficient value.

Robustness profiles for phosphate, thiamine, and riboflavin exhibit predominantly linear behaviour, indicating proportional biomass response to increased import. However, the required quantities differ significantly: thiamine and riboflavin are essential in trace amounts, as evidenced by early inflexion points followed by saturation plateaus.

The oxygen uptake profile reveals a triphasic pattern: a steep initial rise in biomass flux reflecting high sensitivity, followed by a plateau suggesting oxygen saturation, and finally, a decline indicating sub-optimality effects at elevated concentrations. Ferric iron and sulphur display similar biphasic behaviour, excluding the saturation phase. Their initial increase supports growth, confirming their limiting role. However, at higher import rates, a decline in biomass flux suggests toxicity or metabolic imbalance.

Substitutability tests revealed identical profiles for riboflavin and flavin mononucleotide (FMN), indicating functional redundancy in biomass formation. Conversely, the differences observed between oxygen and hydrogen peroxide suggest that, although both can function as electron acceptors, biomass production is more sensitive to oxygen, especially under conditions of low availability.

The analysis of all available carbon and nitrogen sources, clustering based on optimal biomass response identified ten and nineteen behavioural groups, respectively. All carbon sources showed proportional biomass increases with no saturation except for group C1, indicating that most tested conditions were suboptimal (Figure 2.B). Among nitrogen sources, groups N1-N15 exhibited biphasic response curves: an initial steep slope indicating a quite strong sensitivity to nitrogen availability, followed by a shallower slope, reflecting diminishing returns (Figure 2.C). Group N16 reached saturation, while groups N17-N19 showed a subsequent decline in biomass production.

### 3.3. Sensitivity of Specialised Metabolite Production to Biomass Objective Function Perturbations

We further investigated whether perturbations in BOF precursor coefficients affect the model’s capacity to produce selected specialised metabolites related to pathways included in the model. Using FVA, we quantified the maximum production fluxes of ten targeted metabolites or their immediate precursors under both default conditions and after ±50% variation in the stoichiometric coefficients of biomass components (Figure 3). All the minimum values of the flux were equal to 0.

Two distinct response patterns emerged. Adjustments to the stoichiometric coefficients of PROTEIN, RNA, and AAPOOL induced substantial changes in the production potential of penicillin (Figure 3.A), roquefortins (Figure 3.C), and ferrichrome-related compounds (Figure 3.E). In particular, modifying the PROTEIN subsystem yielded maximal flux deviations exceeding 0.5 mmol.gDW⁻¹.h⁻¹ for 6-aminopenicillanic acid, 0.3 mmol.gDW⁻¹.h⁻¹ for isopenicillin N, 0.2 mmol.gDW⁻¹.h⁻¹ for HTD, and 0.1 mmol.gDW⁻¹.h⁻¹ for ferrichromes.

In contrast, the production of precursors for isoepoxydon and yanuthones (Figure 3.B), melanin (Figure 3.D), PR-toxin (Figure 3.F), andrastin (Figure 3.G), as well as averufin and versicolorin (Figure 3.H), remained highly stable, with variations below 10⁻⁵ mmol·gDW⁻¹·h⁻¹. Notably, flux variations on the order of 10⁻⁵ to 10⁻⁷, typically observed in MATLAB-based analyses using the COBRA Toolbox, were not reproduced in the Python implementation with COBRApy. In this case, differences between minimum and maximum fluxes after perturbation were consistently around 10⁻¹⁴, supporting the conclusion that the definition of BOF coefficients does not affect the predicted production of these biosynthetic pathways.

## 4. Discussion

### 4.1. Impacts of Biomass Stoichiometry on Growth Rate Robustness

The definition of the BOF is widely recognised as a critical step in GSMN modelling and must be both qualitatively and quantitatively consistent with the biological characteristics of the organism under study (Bernstein et al., 2021; Feist & Palsson, 2010). While these aspects are extensively documented and rigorously addressed in prokaryotic modelling frameworks, they remain comparatively less developed and pose specific challenges when dealing with more complex or non-canonical model organisms (Chan et al., 2017). In such cases, the scarcity or practical difficulty of acquiring reliable experimental data can represent a major limitation. Filamentous fungi, despite their increasing economic and biotechnological relevance (Garg et al., 2025; Jo et al., 2023; Wilken et al., 2019), exemplify this situation. In this context, the systematic inclusion of sensitivity analyses targeting the quantitative definition of the BOF provides a pragmatic strategy to estimate the extent of growth fluctuations induced by uncertainty in stoichiometric coefficients. Moreover, such analyses help prioritise biomass components that warrant experimental validation and offer an integrated view of the model’s plasticity and robustness. It should nevertheless be borne in mind that these elements constitute a preliminary step towards a global understanding of model behaviour, as only the growth optimum is monitored, and the underlying flux distributions could therefore be radically different (Dikicioglu et al., 2015).

Reactant perturbations of the *i*Prub22 BOF highlights the contribution of individual precursors to the modelled growth phenotype. Reactants whose perturbation led to marked deviations in the optimal biomass flux value are considered critical and thus require careful determination. Conversely, minor effects on growth rate indicate a degree of resilience to such stoichiometric variations. Thus, systematic variations in stoichiometric coefficients provide insight into both the essentiality of specific reactants and their potential role as metabolic bottlenecks.

In principle, increasing the stoichiometric coefficient of a biomass precursor raises its biosynthetic demand, which can reduce predicted growth if the compound is limiting under default conditions. Conversely, decreasing a coefficient may relieve metabolic constraints and enhance growth, unless the precursor fulfils an essential structural or functional role; in this case, growth may become infeasible. However, while such responses may be anticipated in a linear modelling framework, metabolic networks often exhibit pronounced non-linear behaviour. Flux redistribution and the activation of alternative pathways can buffer or amplify stoichiometric perturbations, thereby complicating straightforward interpretation. Nonetheless, simplified sensitivity analyses remain valuable for identifying potential bottlenecks and key control nodes within the network.

By applying ±50% variations to the baseline stoichiometric values of biomass reactants or those of their associated subsystems, the model was deliberately subjected to extreme perturbations to quantify the dependence of predicted growth on individual components. With the exception of two cell wall-related reactants, predicted growth rates remained within ranges typically reported for species of the genus *Penicillium* under comparable experimental conditions (Grijseels et al., 2017). Moreover, increases and decreases in stoichiometric coefficients generally produced symmetrical response profiles. These patterns suggest that, for *P. rubens*, the quantitative precision of biomass stoichiometry , although empirically informed (Agren et al., 2013), may be less critical to model performance than its qualitative composition.

Nevertheless, this analysis identified two critical components in the biomass definition, namely UDP-related compounds. This behaviour is exemplified by UDP-*α*-glucose (CPD-12575) and UDP-*N*-acetyl-*α*-D-glucosamine (UDP-N-ACETYL-D-GLUCOSAMINE), two cell wall precursors (Feofilova, 2010) involved in glycogenesis and glycoprotein/glycolipid biosynthesis, respectively. Alterating their stoichiometric coefficients resulted in suboptimal biomass production. In both cases, increasing the coefficients reduced growth, likely due to elevated metabolic costs, whereas decreasing the coefficients led to a complete loss of biomass production. The impossibility of growth under these conditions suggests a disruption of the steady state, potentially associated with intracellular accumulation of these metabolites. Consequently, these results indicate that the coefficients assigned to UDP-related compounds in *i*Prub22, derived from empirical data in *i*AL1006 (Agren et al., 2013), fall within a narrow range that supports optimal model performance. The observed thresholds emphasise the necessity of maintaining accurate stoichiometric balances, at least for these specific biomass reactants.

### 4.2. Growth Sensitivity to Nutrient Conditions

Fungal growth depends on essential nutrients, including carbon, nitrogen, oxygen, hydrogen, phosphorus, and sulphur, which support energy generation, redox balance, and biosynthetic requirements (Meletiadis et al., 2001; Nielsen, 1992). Carbon and nitrogen can be provided as chemically independent compounds (*e.g.,* glucose, ammonia) or combined, as in amino acids, whereas phosphorus and sulphur are typically assimilated as inorganic phosphate and sulphate, respectively. Filamentous fungi exhibit substantial metabolic versatility and can assimilate a broad spectrum of carbon sources, which can trigger basal metabolism (Aguilar-Pontes et al., 2018; Borin & Oliveira, 2022).

Assessing how simulated growth responds to constraints on nutrient exchange reactions, therefore, serves a dual purpose. First, it enables a systematic inventory of the uptake conditions and combinations effectively represented in the model, thereby providing an indirect evaluation of the accuracy and completeness of environmental modelling. Second, it offers insight into the model’s predictive capacity and its ability to capture metabolic flexibility and adaptive responses to diverse nutritional regimes.

Previous work has demonstrated that *i*Prub22 can reproduce known phenotypes and sustain growth across diverse environmental conditions by utilising a variety of metabolites as carbon and/or nitrogen sources (Nègre et al., 2023). To systematically characterise these nutrient dependencies, we performed robustness analyses by constraining the uptake flux of a single nutrient, forcing its consumption while keeping growth as the sole objective function.

Sensitivity analyses of *i*Prub22 revealed distinct biomass responses to variations in nutrient uptake rates. Linear profiles suggest proportional responses of biomass flux to increasing nutrient availability. Saturated profiles indicate thresholds beyond which further uptake does not enhance growth, whereas inhibitory profiles—characterised by reduced biomass at high uptake rates—may reflect metabolic overload or accumulation effects. Plateaus represent regions of relative insensitivity, while abrupt changes in slope highlight critical sensitivity points. These patterns provide insight into nutrient-specific tolerances and potential bottlenecks.

Qualitatively, most simulated responses seem to align with biological expectations. For instance, inhibitory trends for oxygen uptake could be associated with metabolic stress or disruption (Sies, 2015), and iron uptake profiles may reflect limitations in cofactor availability or intracellular homeostasis (Anderson & Frazer, 2017). Nevertheless, quantitative discrepancies remain, particularly for amino acid use. Amino acids differ in carbon-to-nitrogen ratios, catabolic pathways, and energetic yields. For example, the model predicts lower growth rates for alanine, glutamate, or aspartate than empirically expected, while complex amino acids such as lysine support comparatively higher growth (Nielsen, 1997). These deviations likely reflect limitations in reaction curation, cofactor coupling, or flux partitioning, which require further refinement to improve predictive accuracy.

Conversely, other nutrient simulations align well with experimental evidence. In agreement with the observations of Allam and Elzainy (1969), hypoxanthine and adenine were identified as favourable nitrogen sources, followed by xanthine, urea, and nitrate. These substrates form coherent patterns, reflecting shared assimilation pathways and comparable metabolic efficiencies; thereby supporting the structural consistency of specific sub-networks within *i*Prub22. Specific uptake profiles further suggest the presence of modelling artefacts. Thiamine and riboflavin, which are required only in trace amounts, exhibited linear biomass responses followed by plateaus. Such behaviour could indicate that these compounds may unlock otherwise infeasible cycles (Fritzemeier et al., 2017) within the network rather than representing true growth-limiting nutrients. Similarly, forced uptake, although not physiologically realistic, resulted in reduced biomass production for oxygen, sulphur, phosphate, and nitrogen sources lacking carbon atoms. Interpretation of nitrogen source effects should therefore be approached with caution, as it is not always possible to disentangle whether improved growth arises from nitrogen assimilation itself or from the accompanying carbon contribution.

These simulations were conducted under strictly unitary conditions: the influence of nitrogen sources was evaluated exclusively with glucose as the carbon source, while carbon sources were assessed solely with ammonia as the nitrogen source. Glucose and ammonia were chosen as reference substrates due to their widespread use in fungal culture media. While this design provides direct comparisons, it does not capture potential nutrients interactions. Future combinatorial analyses could provide deeper understanding of metabolic co-dependencies.

Nevertheless, the breadth of the flux ranges delineated by these analyses allows, when required, a more precise definition of which uptake constraints may be relaxed or tightened. In the longer term, such information could provide valuable insight into the rational design and optimisation of culture media. In particular, selecting uptake ranges corresponding to saturation regimes may be most appropriate when the aim is to preserve metabolic capacity for processes decoupled from growth, such as the production of specialised metabolites.

By focusing first on the quantitative definition of the BOF and, subsequently, on the ability of *i*Prub22 to sustain growth across diverse nutritional conditions, these simple analyses provide an integrated overview of the model’s plausible adaptive behaviour. They primarily inform on responses driven by basal metabolism, thereby establishing a foundation for interpreting more complex, condition-specific simulations.

### 4.3. Impacts of Biomass Stoichiometry on Specialised Metabolism

Specialised metabolites, including non-ribosomal peptides, polyketides, and terpenes, confer ecological and competitive advantages to their producing organisms (Rokas et al., 2020), yet their biosynthesis is not intrinsically coupled to growth optimisation (Casas López et al., 2004; Hope et al., 2005). Evaluating how predictions of specialised metabolite production respond to variations in biomass reaction definition provides insight into the stability of modelled phenotypes. It also highlights how biosynthetic capacity is constrained by growth-related demands. Simulations performed with *i*Prub22 showed consistent and biologically plausible responses of specialised metabolic pathways to perturbations in biomass composition.

Perturbations of biomass stoichiometric coefficients did not impose the activation of specialised metabolic pathways, as reflected by minimum flux values consistently equal to zero in flux variability analyses. This indicates that the production of specialised metabolites is either not required or interchangeable to achieve optimal growth. However, changes in BOF composition modulated the maximum production capacities through some specialised metabolism pathways, thereby affecting biosynthetic potential rather than growth requirement. Such a modulation occurred for non-ribosomal peptide pathways (Walsh, 2016), including penicillins, roquefortines, and ferrichromes, whose biosynthesis relies heavily on amino acid availability. Accordingly, the largest shifts in maximum predicted fluxes were observed following perturbations of the PROTEIN and AAPOOL subsystems, which represent the principal mobilisation of amino acids within the model. Modifying these subsystems directly alters the pool of available amino acids for non-ribosomal peptide biosynthesis, thereby constraining or expanding production capacity without altering growth.

In contrast, specialised pathways related to polyketides (*i.e.*, isoepoxydon, yanuthones, melanin, andrastin, averufin, and versicolorin) and terpenes (*i.e.,* PR-toxin) exhibited only marginal sensitivity to biomass perturbations. As these pathways predominantly depend on acetyl-CoA and malonyl-CoA (Nivina et al., 2019; Yin & Dickschat, 2023), also precursors for numerous lipids (Beld et al., 2014; Strijbis & Distel, 2010), their independence from biomass composition suggests that central carbon availability remains largely unaffected by the tested variations. Minor flux changes observed in these pathways likely reflect solver sensitivity or subtle shifts in global flux distributions rather than direct stoichiometric constraints.

Despite the magnitude of the applied stoichiometric perturbations (±50%), variations in specialised metabolite production remained within biologically plausible ranges. This behaviour reflects a stable phenotypic structure in which growth, viability, and the facultative nature of specialised metabolism are preserved, while biosynthetic capacity remains adjustable. Taken together, these results indicate that the quantitative formulation of the BOF exerts limited influence on the activation of specialised metabolic pathways, but modulates their maximal production potential, with non-ribosomal peptides representing the most sensitive class due to their dependence on amino acid metabolism. Overall, the robustness of *i*Prub22 across biomass perturbations supports its suitability for simulating specialised metabolism and identifies amino acid availability as a primary determinant of biosynthetic responsiveness.

## 5. Conclusion

Given that the definition of the biomass reaction is a key determinant of the reliability of GSMNs evaluating whether, and to what extent, flux simulations are influenced by modelling choices for the BOF provides critical insight into overall model quality (Bernstein et al., 2021). The objective of this study is to assess how perturbations on the biomass reaction and environmental conditions affect the overall model predictions. Thus, two complementary strategies were applied (biomass definition perturbation and nutrient flux alteration). Together, these analyses revealed variation patterns that remained within a biologically reasonable range and did not identify any limiting inconsistencies or critical weaknesses in *i*Prub22.

First, we investigated quantitative variations in the biomass reaction and its constituent subsystems to determine whether simulated growth was sensitive to changes in stoichiometric composition. The analysis revealed that the biomass reaction in *i*Prub22 is broadly robust to most stoichiometric perturbations. However, it exhibits pronounced sensitivity to a specific subset of precursors involved in cell wall biosynthesis. These metabolites define narrow tolerance thresholds, beyond which both depletion and excess compromise cellular growth. Accurate experimental calibration of their stoichiometric coefficients is, therefore, critical for maintaining the model’s predictive reliability.

Then, we performed single-reaction robustness analyses on uptake reactions to simulate variations in culture medium composition. This approach allowed us to assess metabolic tolerance to fluctuation of nutrient availability, identify critical thresholds affecting viability, and evaluate metabolic flexibility under alternative nutritional scenarios. Our results confirm that *i*Prub22 captures fundamental metabolic responses to nutrient availability, revealing linear, threshold, and inhibitory relationships that delineate network flexibility and constraints. Nonetheless, the present study is limited to single-variable perturbations. Extending this framework to combinatorial robustness analyses, such as dual-nutrient perturbations or phenotype phase plane analyses (Edwards et al., 2002), would provide deeper insight into nutrient interactions and further enhance the model’s applicability for systems-level predictions.

Finally, among specialised metabolic pathways, only those dependent on non-ribosomal peptide synthase routes exhibited marked sensitivity to biomass formulation. This indicates that predicted specialised metabolite production primarily reflects indirect effects of the biomass objective function on amino acid availability, rather than direct coupling to growth.

Besides highlighting the importance of such sensitivity analysis targeting biomass, a strategy that should be performed systematically during GSMN validation, the new aspect proposed in this manuscript relies on achieving additional sensitivity analysis monitoring reactions of interest before any use of GSMN. In the case presented here, focussing of specialised metabolism of *P. rubens*, it remains important to evaluate related pathways’ sensitivity to the BOF definition. This sensitivity was demonstrated to be directly related to the biosynthetic precursor relation to biomass subsystems. However, exploring such a relation at a very large scale across GSMNs remains a challenging task as specialised metabolism is intrinsically species-specific, and even strain-specific (Wei et al., 2025). Extracting dedicated pathways needs a careful examination of the literature and specific genome mining tools (Blin et al., 2025; Wang et al., 2025). For such goals, additional pathway labelling should be present in databases such as MetaCyc (Karp et al., 2019) and KEGG (Kanehisa et al., 2025), and a consistent definition should be provided (Nègre et al., 2023). Through these examples, we aimed to underline the relevance of simple, rapid, and pre-existing analyses for GSMN evaluation and refinement. Despite their low computational cost and straightforward implementation, such tests remain, in our opinion, predominantly underused in model reconstruction workflows. Their systematic inclusion would enhance the transparency, reliability, and biological interpretability of published GSMNs, supporting both their validation and broader applicability in predictive systems biology.

However, it should be noted that this study focused on the quantitative definition of the biomass reaction rather than its qualitative composition, an essential yet frequently overlooked aspect of GSMN reconstruction (Bernstein et al., 2021). This limitation is less critical for *P. rubens*, as relevant data addressing generic biomass composition are available (Agren et al., 2013; Nielsen, 1997; Van den Berg et al., 2008). Nonetheless, compared with the earlier model *i*AL1006 (Agren et al., 2013), the updated reconstruction *i*Prub22 lacks several metabolic subsystems, notably those involved in lipid biosynthesis (*i.e.,* fatty acids, neutral lipids, and glycerides). Lipid metabolism remains a particularly challenging domain for reconstruction despite recent efforts to improve annotation and ontology coverage (Poupin et al., 2020).

Indeed, beyond the definition of the BOF, flux distributions within a GSMN are primarily constrained by network topology, a factor of particular importance given the partial representation of specialised metabolism in current reconstruction databases (Qiu et al., 2023). When present, specialised metabolic pathways in GSMNs derived from semi-automatic or fully manual processes often reflect user-driven, partially arbitrary choices, similar to those underlying biomass objective function formulation. Nevertheless, for *i*Prub22, a GSMN describing a multicellular eukaryote, the results presented here indicate a solid and generic modelling framework. This framework provides a consistent basis for subsequent refinements, including the formulation of alternative biomass objective functions tailored to specific nutritional environments or distinct cellular types, thereby supporting future context-dependent modelling applications (Stalidzans & Dace, 2021).

## 6. Data availability statement

The model used in this study is the GSMN *i*Prub22 of *Penicillium rubens*, publicly accessible *via* BioModels (MODEL2306150001). Computational workflows (*i.e.,* scripts and result files) required to reproduce all analyses are archived on Zenodo (https://doi.org/10.5281/zenodo.17091974).

---

**Table S1.**
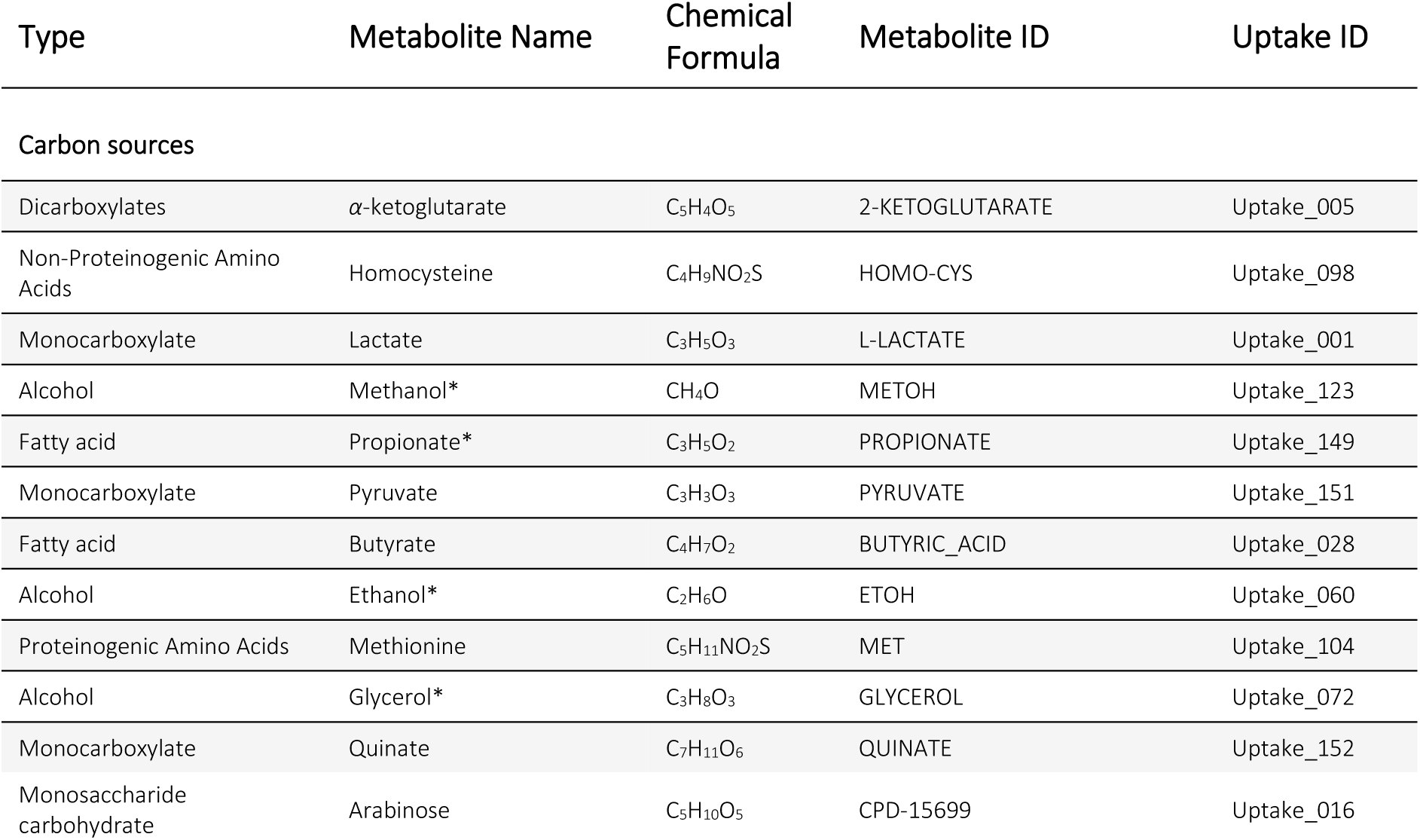
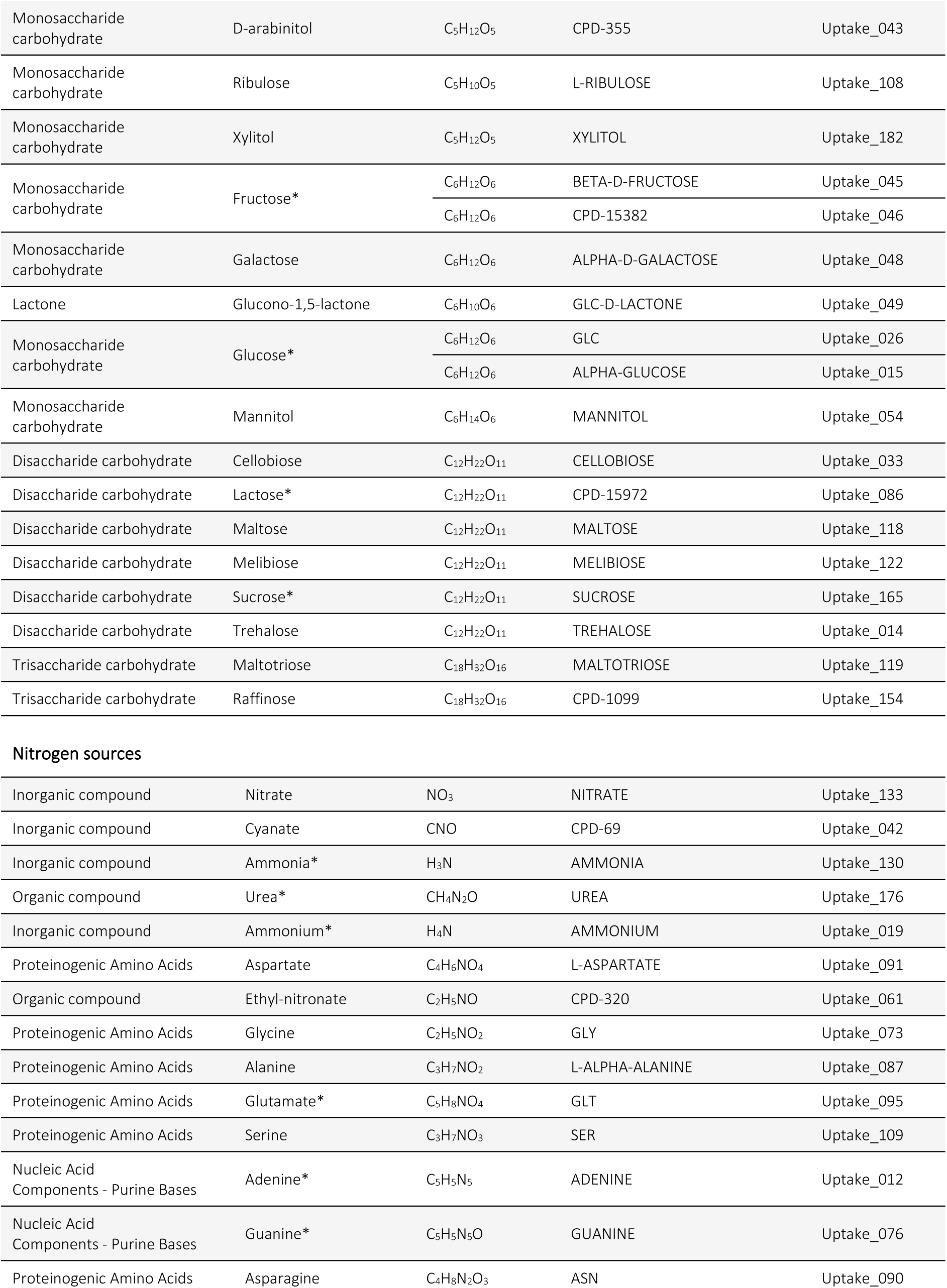
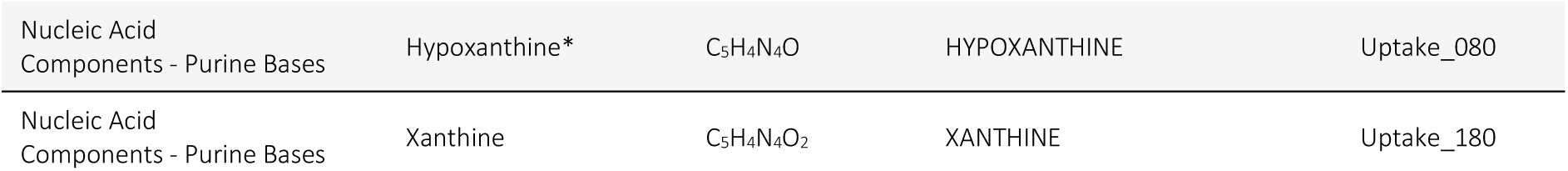
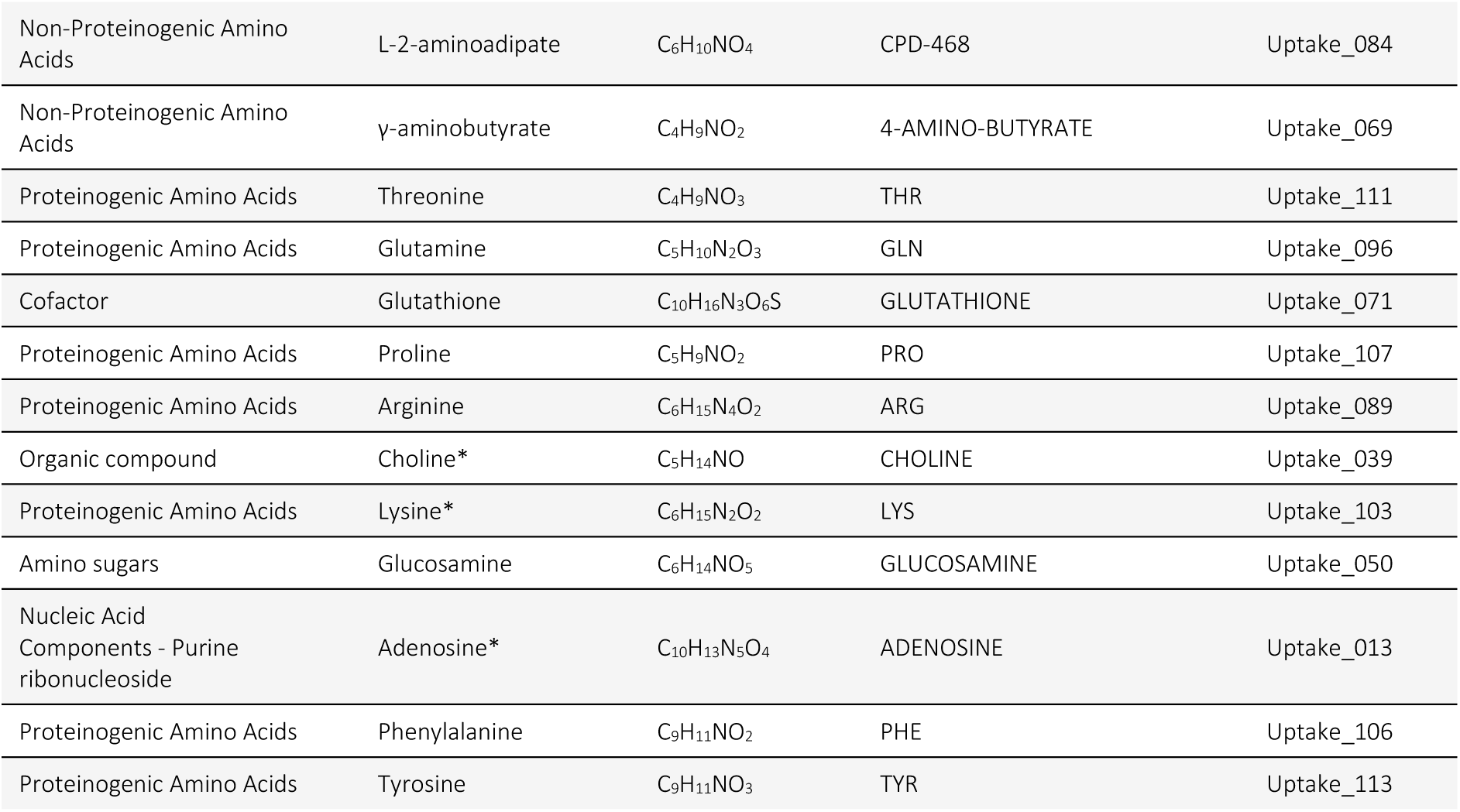
Metabolites supporting growth as sole carbon or nitrogen sources in iPrub22. This table presents metabolites tested individually as sole carbon or nitrogen sources under defined minimal conditions, while maintaining the iPrub22 model functionality. Model constraints were consistent with those determined by default. Only compounds that sustain a non-zero biomass flux are included. Carbon and nitrogen sources are categorised accordingly and identified by their compound name and corresponding uptake reaction. Elements are presented in decreasing order of their potential efficiency in supporting biomass production in iPrub22 simulations. Terms in bold followed by an asterisk (*) are derived from published methodologies, such as the patent on the fermentative production of industrially relevant compounds using chemically defined media (Laat et al., 2002).

| Type | Metabolite Name | Chemical Formula | Metabolite ID | Uptake ID |
| --- | --- | --- | --- | --- |
| **Carbon sources** |  |  |  |  |
| Dicarboxylates | α-ketoglutarate | C<sub>5</sub>H<sub>4</sub>O<sub>5</sub> | 2-KETOGLUTARATE | Uptake_005 |
| Non-Proteinogenic Amino Acids | Homocysteine | C<sub>4</sub>H<sub>9</sub>NO<sub>2</sub>S | HOMO-CYS | Uptake_098 |
| Monocarboxylate | Lactate | C<sub>3</sub>H<sub>5</sub>O<sub>3</sub> | L-LACTATE | Uptake_001 |
| Alcohol | Methanol* | CH<sub>4</sub>O | METOH | Uptake_123 |
| Fatty acid | Propionate* | C<sub>3</sub>H<sub>5</sub>O<sub>2</sub> | PROPIONATE | Uptake_149 |
| Monocarboxylate | Pyruvate | C<sub>3</sub>H<sub>3</sub>O<sub>3</sub> | PYRUVATE | Uptake_151 |
| Fatty acid | Butyrate | C<sub>4</sub>H<sub>7</sub>O<sub>2</sub> | BUTYRIC_ACID | Uptake_028 |
| Alcohol | Ethanol* | C<sub>2</sub>H<sub>6</sub>O | ETOH | Uptake_060 |
| Proteinogenic Amino Acids | Methionine | C<sub>5</sub>H<sub>11</sub>NO<sub>2</sub>S | MET | Uptake_104 |
| Alcohol | Glycerol* | C<sub>3</sub>H<sub>8</sub>O<sub>3</sub> | GLYCEROL | Uptake_072 |
| Monocarboxylate | Quinate | C<sub>7</sub>H<sub>11</sub>O<sub>6</sub> | QUINATE | Uptake_152 |
| Monosaccharide carbohydrate | Arabinose | C<sub>5</sub>H<sub>10</sub>O<sub>5</sub> | CPD-15699 | Uptake_016 |

**Table S1 (continued)**
|  |  |  |  |  |
| --- | --- | --- | --- | --- |
| Monosaccharide carbohydrate | D-arabinitol | C<sub>5</sub>H<sub>12</sub>O<sub>5</sub> | CPD-355 | Uptake_043 |
| Monosaccharide carbohydrate | Ribulose | C<sub>5</sub>H<sub>10</sub>O<sub>5</sub> | L-RIBULOSE | Uptake_108 |
| Monosaccharide carbohydrate | Xylitol | C<sub>5</sub>H<sub>12</sub>O<sub>5</sub> | XYLITOL | Uptake_182 |
| Monosaccharide carbohydrate | Fructose* | C<sub>6</sub>H<sub>12</sub>O<sub>6</sub> | BETA-D-FRUCTOSE | Uptake_045 |
|  |  | C<sub>6</sub>H<sub>12</sub>O<sub>6</sub> | CPD-15382 | Uptake_046 |
| Monosaccharide carbohydrate | Galactose | C<sub>6</sub>H<sub>12</sub>O<sub>6</sub> | ALPHA-D-GALACTOSE | Uptake_048 |
| Lactone | Glucono-1,5-lactone | C<sub>6</sub>H<sub>10</sub>O<sub>6</sub> | GLC-D-LACTONE | Uptake_049 |
| Monosaccharide carbohydrate | Glucose* | C<sub>6</sub>H<sub>12</sub>O<sub>6</sub> | GLC | Uptake_026 |
|  |  | C<sub>6</sub>H<sub>12</sub>O<sub>6</sub> | ALPHA-GLUCOSE | Uptake_015 |
| Monosaccharide carbohydrate | Mannitol | C<sub>6</sub>H<sub>14</sub>O<sub>6</sub> | MANNITOL | Uptake_054 |
| Disaccharide carbohydrate | Cellobiose | C<sub>12</sub>H<sub>22</sub>O<sub>11</sub> | CELLOBIOSE | Uptake_033 |
| Disaccharide carbohydrate | Lactose* | C<sub>12</sub>H<sub>22</sub>O<sub>11</sub> | CPD-15972 | Uptake_086 |
| Disaccharide carbohydrate | Maltose | C<sub>12</sub>H<sub>22</sub>O<sub>11</sub> | MALTOSE | Uptake_118 |
| Disaccharide carbohydrate | Melibiose | C<sub>12</sub>H<sub>22</sub>O<sub>11</sub> | MELIBIOSE | Uptake_122 |
| Disaccharide carbohydrate | Sucrose* | C<sub>12</sub>H<sub>22</sub>O<sub>11</sub> | SUCROSE | Uptake_165 |
| Disaccharide carbohydrate | Trehalose | C<sub>12</sub>H<sub>22</sub>O<sub>11</sub> | TREHALOSE | Uptake_014 |
| Trisaccharide carbohydrate | Maltotriose | C<sub>18</sub>H<sub>32</sub>O<sub>16</sub> | MALTOTRIOSE | Uptake_119 |
| Trisaccharide carbohydrate | Raffinose | C<sub>18</sub>H<sub>32</sub>O<sub>16</sub> | CPD-1099 | Uptake_154 |
| Nitrogen sources |  |  |  |  |
| Inorganic compound | Nitrate | NO<sub>3</sub> | NITRATE | Uptake_133 |
| Inorganic compound | Cyanate | CNO | CPD-69 | Uptake_042 |
| Inorganic compound | Ammonia* | H<sub>3</sub>N | AMMONIA | Uptake_130 |
| Organic compound | Urea* | CH<sub>4</sub>N<sub>2</sub>O | UREA | Uptake_176 |
| Inorganic compound | Ammonium* | H<sub>4</sub>N | AMMONIUM | Uptake_019 |
| Proteinogenic Amino Acids | Aspartate | C<sub>4</sub>H<sub>6</sub>NO<sub>4</sub> | L-ASPARTATE | Uptake_091 |
| Organic compound | Ethyl-nitronate | C<sub>2</sub>H<sub>5</sub>NO | CPD-320 | Uptake_061 |
| Proteinogenic Amino Acids | Glycine | C<sub>2</sub>H<sub>5</sub>NO<sub>2</sub> | GLY | Uptake_073 |
| Proteinogenic Amino Acids | Alanine | C<sub>3</sub>H<sub>7</sub>NO<sub>2</sub> | L-ALPHA-ALANINE | Uptake_087 |
| Proteinogenic Amino Acids | Glutamate* | C<sub>5</sub>H<sub>8</sub>NO<sub>4</sub> | GLT | Uptake_095 |
| Proteinogenic Amino Acids | Serine | C<sub>3</sub>H<sub>7</sub>NO<sub>3</sub> | SER | Uptake_109 |
| Nucleic Acid Components - Purine Bases | Adenine* | C<sub>5</sub>H<sub>5</sub>N<sub>5</sub> | ADENINE | Uptake_012 |
| Nucleic Acid Components - Purine Bases | Guanine* | C<sub>5</sub>H<sub>5</sub>N<sub>5</sub>O | GUANINE | Uptake_076 |
| Proteinogenic Amino Acids | Asparagine | C<sub>4</sub>H<sub>8</sub>N<sub>2</sub>O<sub>3</sub> | ASN | Uptake_090 |
| Nucleic Acid Components - Purine Bases | Hypoxanthine* | C<sub>5</sub>H<sub>4</sub>N<sub>4</sub>O | HYPOXANTHINE | Uptake_080 |
| Nucleic Acid Components - Purine Bases | Xanthine | C<sub>5</sub>H<sub>4</sub>N<sub>4</sub>O<sub>2</sub> | XANTHINE | Uptake_180 |

**Table S1 (continued)**
|  |  |  |  |  |
| --- | --- | --- | --- | --- |
| Non-Proteinogenic Amino Acids | L-2-aminoadipate | C<sub>6</sub>H<sub>10</sub>NO<sub>4</sub> | CPD-468 | Uptake_084 |
| Non-Proteinogenic Amino Acids | γ-aminobutyrate | C<sub>4</sub>H<sub>9</sub>NO<sub>2</sub> | 4-AMINO-BUTYRATE | Uptake_069 |
| Proteinogenic Amino Acids | Threonine | C<sub>4</sub>H<sub>9</sub>NO<sub>3</sub> | THR | Uptake_111 |
| Proteinogenic Amino Acids | Glutamine | C<sub>5</sub>H<sub>10</sub>N<sub>2</sub>O<sub>3</sub> | GLN | Uptake_096 |
| Cofactor | Glutathione | C<sub>10</sub>H<sub>16</sub>N<sub>3</sub>O<sub>6</sub>S | GLUTATHIONE | Uptake_071 |
| Proteinogenic Amino Acids | Proline | C<sub>5</sub>H<sub>9</sub>NO<sub>2</sub> | PRO | Uptake_107 |
| Proteinogenic Amino Acids | Arginine | C<sub>6</sub>H<sub>15</sub>N<sub>4</sub>O<sub>2</sub> | ARG | Uptake_089 |
| Organic compound | Choline* | C<sub>5</sub>H<sub>14</sub>NO | CHOLINE | Uptake_039 |
| Proteinogenic Amino Acids | Lysine* | C<sub>6</sub>H<sub>15</sub>N<sub>2</sub>O<sub>2</sub> | LYS | Uptake_103 |
| Amino sugars | Glucosamine | C<sub>6</sub>H<sub>14</sub>NO<sub>5</sub> | GLUCOSAMINE | Uptake_050 |
| Nucleic Acid Components - Purine ribonucleoside | Adenosine* | C<sub>10</sub>H<sub>13</sub>N<sub>5</sub>O<sub>4</sub> | ADENOSINE | Uptake_013 |
| Proteinogenic Amino Acids | Phenylalanine | C<sub>9</sub>H<sub>11</sub>NO<sub>2</sub> | PHE | Uptake_106 |
| Proteinogenic Amino Acids | Tyrosine | C<sub>9</sub>H<sub>11</sub>NO<sub>3</sub> | TYR | Uptake_113 |

